# The Association of Externalizing and Internalizing Problems with Indicators of Intelligence in a Sample of At-Risk Children

**DOI:** 10.1101/210500

**Authors:** Nicholas Kavish, Jesse Helton, Michael G. Vaughn, Brian B. Boutwell

## Abstract

To date, a substantial body of research exists suggesting an association between indicators of intelligence and various deleterious outcomes, including externalizing and internalizing behavioral problems. Much of this research, however, has focused on samples drawn from the general population, thus it remains less clear how (and if) intelligence relates to problem behaviors in samples of highly at-risk individuals. The current study seeks to contribute to this knowledge base by examining the associations between intelligence and internalizing, externalizing, and total scores on the Child Behavioral Checklist in a sample of approximately 2,500 highly disadvantaged respondents considered by Child Protective Services as at-risk for abuse or neglect. While the two measures of intelligence performed differently, there emerged some association between overall lower IQ and higher total behavioral problem scores. There was some evidence that low IQ also predicted higher internalizing scores, but this relationship varied greatly by measure and model. Results, limitations, and implications of the current study are discussed.

## Introduction

The study of human intelligence over the last century has continued to provide meaningful insights about a range of important developmental and social outcomes (Kline, 2013; Ritchie, 2015). Indeed, what seems rather clear at this point, is that variation in intelligence predicts variation across key social outcomes across all phases of the life course, beginning in childhood and spanning into adulthood (Gottfredson, 2004; Plomin & Deary, 2015; Ritchie, 2015). In what constitutes one of the more notable tests on the topic, Moffitt and colleagues (2011), using a sample of respondents tracked from birth until the third decade of life, uncovered evidence that along with self-control, intelligence was consistently associated with indicators of health and economic success across years of the lifespan.

Other studies have uncovered very similar patterns of effects, revealing positive associations between intelligence and educational attainment, accrual of wealth, increased self-regulation, upward social mobility, selection of friends and romantic partners (i.e., assortative paring), and even longer life expectancy, across multiple independent samples (Arden et al., 2015; Beaver et al., 2016; Calvin et al., 2017; Gottfredson, 2004; Boutwell et al., 2017; Meldrum et al., 2017; Plomin and Deary, 2015). Not only have higher levels of intelligence been linked to positive social outcomes, the inverse also seems to be true, in that lower levels of intelligence predict various adverse life events. Beaver et al (2016), for instance, using a nationally representative sample of over 15,000 participants, found that those in the bottom 25% of IQ scores were almost twice as likely to be victimized as those in the top 25%. Lower levels of intelligence are also associated with an increased risk of self-reported criminal justice processing (being arrested and incarcerated) in adulthood (Beaver et al., 2013).

Finally, Calvin and colleagues (2017) recently analyzed a large sample of Scottish participants, and uncovered an association between childhood intelligence and various causes of mortality across several decades of the life course. In particular, children who scored higher on childhood measures of intelligence, were less likely to die from a variety of adulthood conditions such as coronary heart disease and smoking related cancer (see Calvin et al., 2017 for additional detail; see also, Arden et al., 2015). Though far from exhaustive, this body of evidence clearly suggests that variation in measures of intelligence represents a robust correlate for a wide swath of phenotypes (see also Aarons, James, Monn, Raghavan, Wells, & Leslie, 2010; Ritchie, 2015; Vaughn, Shook, & McMillen, 2008).

### Remaining Questions to Ask About Intelligence

Despite the consistent pattern of findings briefly outlined above, there remains several interesting gaps in the literature that need to be examined. In particular, less effort to date has been aimed at examining whether, and to what extent, intelligence predicts adverse or antisocial outcomes in samples of highly at-risk participants. Children exposed to abuse, trauma, neglect, and maltreatment on the part of their caregivers, for instance, are at risk for a host of maladaptive outcomes, psychological and behavioral problems among them (Flouri, Midouhas, & Joshi, 2015; Jonson-Reid & Drake, 2017). Given the broad nexus of risk factors these children are often exposed to (in general, see Jonson-Reid & Drake, 2017), it seems important to further explore whether indicators of intelligence might explain variation for behavioral and psychological problems in a population broadly exposed to a range of risk factors.

Additionally, most of the studies outlined above examined openly manifested forms of antisocial behavior (i.e., violence, arrests, etc.), as opposed to using more clinically relevant instruments such as scales designed to assess externalizing psychopathology early in the life course. Even less work (relatively speaking), has specifically focused on internalizing psychopathology. Furthermore, very little evidence exists concerning the association between intelligence and *both* externalizing and internalizing problems in a sample of extremely disadvantaged respondents—such as children exposed to abuse and neglect early in life. Below we discuss some of the literature that has attempted to address this topic, then transition to outlining the goals of the current study.

### Intelligence and Psychopathology

As a brief reminder, psychopathology—from a clinical standpoint—can be broadly subsumed in two main categories: externalizing and internalizing problems (Caspi et al., 2014; Kotov et al., 2017), with two smaller categories sometimes reported for thought disorder/psychotic experiences and for somatic symptoms depending on the sample (Marek et al., 2019; Wright et al., 2013). These main hierarchical clusters (internalizing and externalizing) have been replicated cross-culturally (Kessler et al., 2011) and across the life course, manifesting in both children/adolescents (Achenbach, 1966; Lahey et al, 2011) and adults (Krueger & Markon, 2006). Generally speaking, externalizing problems encompass overt displays of aggression and impulsivity (del Giudice, 2016). Internalizing problems, on the other hand, involve difficulties with anxiety, depression, and other less overt forms of psychopathology. Researchers examining the origins and nature of psychopathology have long recognized the tendency for externalizing and internalizing problems to be comorbid with one another (Caspi et al., 2014; Lilienfeld, 2003), strongly suggesting a shared vulnerability across domains (often referred to as the “p-factor”; Caspi et al., 2014). Put another way, individuals with internalizing (or externalizing) problems, are at higher odds of experiencing some form of the other domain of psychopathology.

At the same time, extant research has examined the possible risk factors, both phenotypic and genetic, which might account for variation within these domains of symptoms, and also perhaps shed light on the co-occurrence of internalizing and externalizing problems (Del Giudice, 2016). To date, intelligence seems to have a clear association with externalizing problems (Guay, Ouimet, & Proulx, 2005; Hinshaw, 1992; Menting, Hirschi & Hindelang, 1977; Menting, Van Lier & Koot, 2011; see also Aarons, James, Monn, Raghavan, Wells, & Leslie, 2010; Vaughn, Shook, & McMillen, 2008). In a review of the longitudinal research on IQ, school achievement, and externalizing behavior problems, Hinshaw (1992) concluded that lower IQ was a strong predictor of increased behavioral problems. Duran-Bonavlila and colleagues (2017) found intelligence was negatively correlated with physical, indirect (e.g., gossiping, socially ostracizing others), and overall aggression in a sample of Spanish high school students, even after controlling for impulsivity. Low cognitive ability has also been *causally* associated with increased risk for alcohol use disorders using genetically sensitive analyses in a sample of around 1 million Swedes (Kendler et al., 2017). Finally, intelligence has been found to be negatively, albeit weakly, with measures assessing conduct disorder, antisocial personality disorder, and psychopathic traits (Sánchez de Ribera et al., 2019).

Less research to date has examined the association between intelligence and internalizing difficulties. One study found verbal ability (but not math ability) at ages 4-5 was negatively related to self-reported internalizing symptoms at ages 12-15 (Weeks et al., 2015). Wraw and colleagues (2016) examined IQ at age 15-23 and mental health at age 50 in a sample of over 5,000 participants and found that a lower IQ was related to higher self-reported depression symptoms and poorer self-reported overall mental health (but paradoxically lower odds of reporting a lifetime diagnosis of depression). Similarly, Gale et al. (2008) found that lower cognitive ability was related in increased risk for depression, generalized anxiety, and PTSD in a prospective cohort study of Vietnam veterans.

### Intelligence and Psychopathology in Disadvantaged Populations

Despite the relatively clear evidence that lower intelligence is associated with increased risk for psychopathology, especially externalizing difficulties, across a number of populations, it remains unclear whether similar findings would emerge in highly disadvantaged and at-risk samples of children —children, for example, who are members of families that have been actively investigated by Child Protective Services (CPS) for alleged abuse or neglect. One recent study has helped to shed some insight on to this topic, albeit with a broad focus on adversity, rather than on alleged abuse or neglect. Using a large sample including overrepresentation of socioeconomically disadvantaged families in the United Kingdom, Flouri, Midouhas, & Joshi (2015) found that intelligence measured between ages 3 and 7 was associated with *both* internalizing and externalizing disorders. Interestingly, the evidence that intelligence moderates the relationship between environmental threats (socioeconomic disadvantage, adverse life events, and neighborhood poverty) and changes in child externalizing behavioral problems over time was more mixed. Nonetheless, less intelligent children, irrespective of environmental threat, experienced more internalizing and externalizing problems over time compared to more intelligent children.

Additionally, another recent study using a U.S. based group of subjects examined the relationship between measures of intelligence and psychopathology in a longitudinal sample of allegedly maltreated children (Harpur, Polek, & Harmelen, 2015)^1^. The authors found that both higher spatial and verbal intelligence measured at age 6 was associated with lower levels of anxiety and depression at age 14. Thus, preliminary evidence suggests that intelligence is inversely related to both externalizing and internalizing problems across childhood and into early adolescence for children at risk for abuse and neglect, mirroring findings in the general population. Nonetheless more evidence remains required.

## The Current Study

To further examine the association between indicators of intelligence and problem behaviors, the current study makes use of a unique dataset of highly at-risk American children. In doing so, our study benefits from several strengths. First, relative to Harpur and colleages (2015) – the most directly related study on the topic of IQ and psychopathology in children at risk for abuse and neglect – we analyze a larger nationally representative sample of children who have been investigated by CPS based on allegations of abuse and neglect on the part of their caregivers within the United States. The sample represents children who may be considered the most-imperiled members of the population in that they are also predominately minority respondents residing in lower socioeconomic strata. Furthermore, we evaluate the association between intelligence and psychopathology using subtests from two different cognitive tests and have data on internalizing and externalizing difficulties at baseline and follow-up to better assess change over time. To date, it remains less clear whether variation in measures of intelligence are associated (and to what extent) with behavioral problems in this segment of the population. Although this population may have on average more externalizing and internalizing difficulties, we hypothesize an inverse relationship between intelligence and both externalizing and internalizing symptoms: as intelligence scores increase, internalizing and externalizing symptoms should decrease.

## Methods

### Sample

For the current analyses, we employed data drawn from the second cohort of the National Survey of Child and Adolescent Well-Being (NSCAW II). NSCAW II, has been discussed in detail in a variety of other studies and as a result, we restrict our description to an abbreviated discussion. The study designers utilized a two-stage stratified sampling design. The first step was to select nine sampling strata consisting of the eight states with the largest child welfare caseloads and the remainder of the US (see Dolan, Smith, Casanueva, & Ringeisen, 2011). The primary sampling units (PSUs), were then selected within each of the nine strata. The same numbers of families were then sampled within each of the 83 selected PSUs. For the NSCAW II sample, cases from CPS investigations that were closed between February 2008 and April 2009 nationwide (*n* = 5873) were included. The final sample of children was representative of the national population of children birth to 17 years of age in families being investigated for allegations of maltreatment (Dowd et al., 2011).

An important aspect of the sample is that it included both substantiated and unsubstantiated investigations. Moreover, NSCAW II also contained cases that received family preservation services, as well as those who did not receive services in the wake of an investigation. Finally, the sample also included families who had their children removed to foster care, following CPS investigations. Face-to-face interviews with children and current caregiver by trained NSCAW practitioners were completed on average 4 months after the close of the investigation, and again one year (approximately) following the baseline interview (Waves 1 and 2 of NSCAW). Due to age restrictions on our standardized measures, we restricted the data to children over the age of 3 at baseline interview (n=2591).

### Measures

#### Child behavior

The caregiver reported Child Behavioral Checklist (CBCL; Achenbach, 1991) was used to assess behavioral problems in the current study. The CBCL measures internalizing and externalizing symptoms as well as other domains of behavior, including social, thought, and attention problems. For internalizing, three subscales were combined and averaged: somatic complaints, anxiety and depression, and withdrawn. For externalizing problems, two subscales were combined and averaged: rule breaking and aggressive behaviors. A total score was also derived using all of the above subscales with an additional 33 items subsumed under “other problems” domain. The CBCL gives a standardized score with a mean of 50 and standard deviation of 10, with higher scores indicating greater behavioral problems.

#### Indicators of Intelligence

We utilize two indicators of intelligence in the current analysis (see Table 1 for a listing of intelligence test subscales, descriptions, and abilities being assessed). First, the Kaufman Brief Intelligence Test (KBIT; Kaufman & Kaufman, 1990) test was used, which assesses both verbal intelligence (word knowledge and verbal concept formation) and nonverbal intelligence. Additionally, we utilized a normalized sum of standard scores for vocabulary and matrices scores at points in the statistical modeling. The KBIT was administered to children directly by an NSCAW interviewer. Scores ranged from 40 to 142, with a mean of 100, a standard deviation of 15 (in the general population). Second, we examined scores on the Woodcock Johnson III Tests of Achievement (WJ-III). Importantly, correlations between cognitive test score composites and achievement test score composites, including the WJ-III, are typically moderately strong (*r* ∼ .70; Naglieri & Bornstein, 2003). Each scale in the measure— Letter Identification, Passage Comprehension and Applied Problems—has national norms for each age and an average score of 100 and standard deviation of 15 in the general population.

**Table 1.** Cognitive ability measure subtests, descriptions, and the abilities they measure

| **Scale/Subtest** | **Subtest Description** | **Ability measured by the subtest** |
| --- | --- | --- |
| **K-BIT** |  |  |
| Verbal | Verbal knowledge and “riddle” items assessing knowledge of words and their meanings – no reading required | Expressive and receptive vocabulary |
| Matrices | Uses pictures and abstract designs to assess ability to perceive patterns and solve analogies | Non-verbal, fluid reasoning ability |
| **WJ-III-ACH** |  |  |
| Letter Identification | Subject names presented letters or reads presented words | Reading decoding |
| Passage Comprehension | Requires identifying a missing key word that fits based on context of a written passage | Reading comprehension |
| Applied Problems | Requires solving orally presented math problems | Quantitative reasoning, math knowledge |

An important note to mention is that children over the age of 11 (i.e., 12 and over) were not administered the Passage Comprehension tests in NSCAW. Thus, when creating the total WJ-III score for the current study, a child’s average score of the available scale based on age was used. In other words, for the creation of the total WJ-III composite measure, we summed and averaged the scores on available subscales. If the child was 12 years of age or older, and thus were not administered the Passage Comprehension subtest, only the other two subscales were included. If a child was 11 years old or under, all three subscales are used. Finally, we created a summated and averaged total intelligence score based on the total K-BIT and the total WJ-III. To clarify further, every participant possessed a measure of total intelligence, however, children over the age of 11 did not have the Passage Comprehension subscale included in their total intelligence score (given that they were not administered this particular subscale by NSCAW staff; Dowd, Kinsey, Wheeless, et al., 2004).

#### Child demographics

The child’s age, race/ethnicity, and sex were assessed by structured interview with primary caregiver at time of NSCAW interview.

#### Family Poverty

The NSCAW team created a poverty variable that was measured by calculating the family’s income-to-needs ratio, which was estimated by dividing family income by its corresponding poverty threshold in 2009 (Dowd et al, 2004). The poverty threshold varies by family size and is based on the money necessary for the minimally accepted amounts of food, with 1.00 representing the overall poverty threshold (Bishaw & Iceland, 2003). Caregivers reported both family income and household size. This measure was divided into four categories: at or below 50% of the poverty line, between 51% and 100% of the poverty line, between 101% of the poverty line and 200%, and above 200% of the poverty line.

### Analytic Approach

All analyses were performed using STATA Statistical Software Release 13 (Stata, 2013). Due to NSCAW’s complex sampling design, special STATA survey commands were applied to obtain unbiased estimates of population parameters (NSCAW Research Group, 2002). All percentages were weighted for sample probabilities; therefore, percentages reported in tables represent national estimates. To accommodate certain aspects of the survey design, χ^2^ statistics were converted to *F*-statistics with noninteger degrees of freedom using a second-order Rao and Scott correction (StataCorp, 2003). All intelligence and behavioral measures were normally distributed, making ordinary least squares (OLS) regression appropriate. Three separate regressions were used to examine the relationship between intelligence and future behavior. The first regression modeled only behavior at wave 2 by intelligence at wave 1 and the second model added important covariates such as child age, race, sex, and family poverty. In the third model, we added behavior scores at wave 1 as a type of lag variable, thereby providing the ability to test change in behavioral scores across waves as a function of intelligence while still controlling for covariates from baseline (Cohen, Cohen, West, Aiken, 2003). Unstandardized coefficients are presented in tables. Missing data were limited to 40 cases, or 1.5 percent of the sample.

## Results

Table 2 reports a description of the sample. Average intelligence scores in the sample were lower than the population mean (mean score is typically 100). Average child age was 10 years, gender was equally distributed, and less than half of the sample was non-Hispanic white. Over half (54.7%) of children were living in families that were under the federal poverty line. Twenty-four percent of CPS investigations were found credible (i.e. substantiated), and 60% of children remained in-home without CPS services, 24% remained in-home with CPS services, and 16% were removed from care and placed with an out-of-home caregiver (results not reported in tables).

**Table 2.**
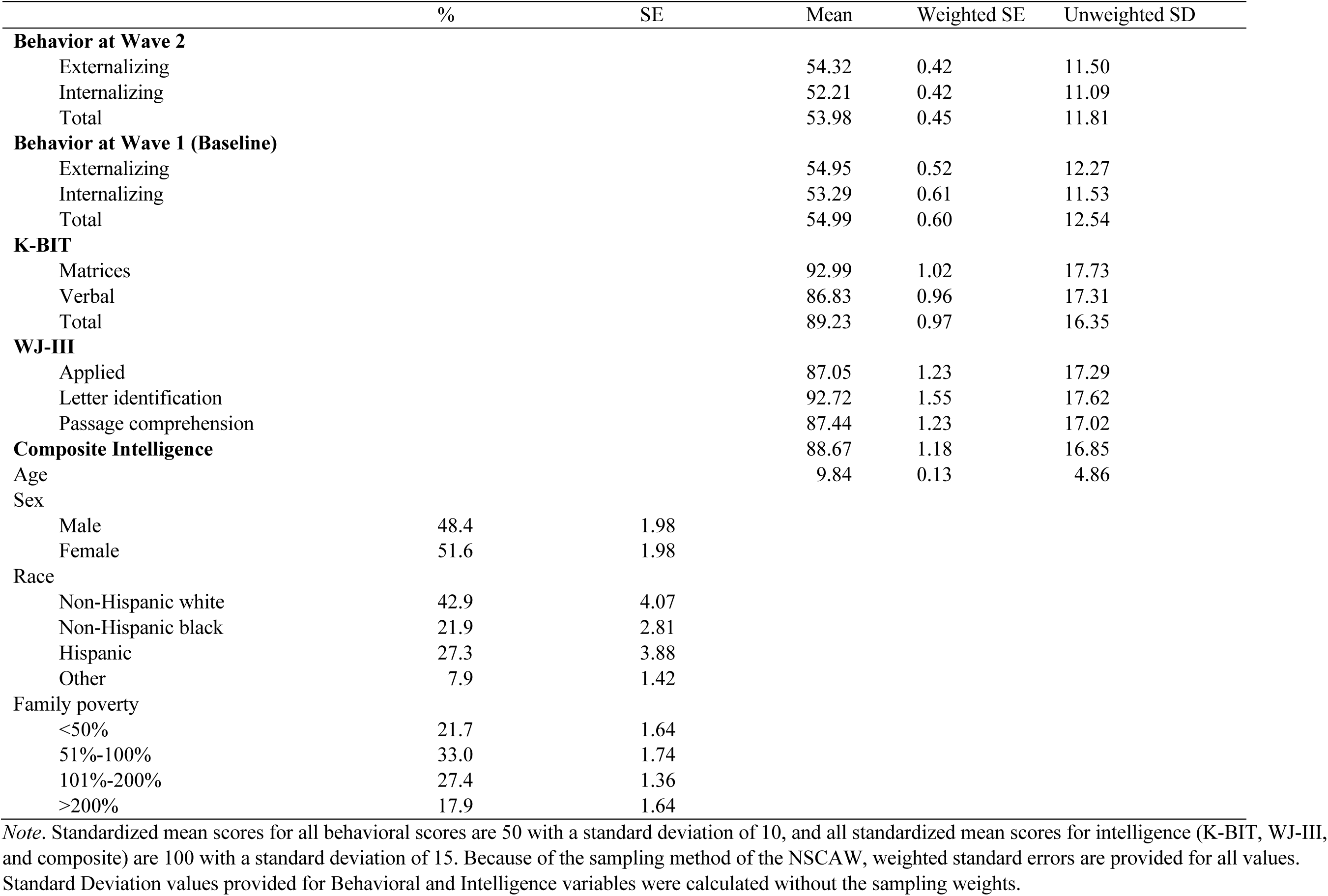
Sample characteristics (n=2591)

### Does intelligence at baseline predict psychopathology measured one year later?

Tables 3 through 5 present results of regression models predicting externalizing (Table 3), internalizing (Table 4) and total behavior problems (Table 5). Across these tables, model one explores the question of whether or not there is an association between intelligence indicators and the relevant psychopathology outcome. For externalizing, most intelligence variables were inversely related to symptom severity. That is, as scores on all intelligence variables increased – with the exception of Verbal on the K-BIT and Passage Comprehension on the WJ-III – parent ratings of externalizing symptoms decreased. Conversely, intelligence was generally unrelated to internalizing symptoms one year later, with the exception of the WJ-III Letter Identification subtest and total scores, which were negatively related to symptomology. Finally, results were mixed with regard to total problems, as the K-BIT and its subtests were not statistically related to parent ratings, but WJ-III Applied Problems, Letter Identification, and total score, as well as the intelligence composite, were all inversely related to overall behavior problems. In sum, intelligence appears consistently, negatively related to externalizing difficulties, mostly unrelated to internalizing symptoms, and inconsistently, negatively related to overall behavior problems as rated by parents one year later.

**Table 3:** Regression models predicting externalizing CBCL conditions at time 2

|  | Model 1 |  | Model 2 |  | Model 3 |  |
| --- | --- | --- | --- | --- | --- | --- |
|  | *b* | *SE* | *b* | *SE* | *b* | *SE* |
| **K-BIT** |  |  |  |  |  |  |
| Matrices | -0.06* | 0.03 | -0.04 | 0.03 | -0.06** | 0.02 |
| Externalizing (Baseline) |  |  |  |  | 0.62** | 0.03 |
| Verbal | -0.04 | 0.02 | -0.04 | 0.03 | -0.04* | 0.02 |
| Externalizing (Baseline) |  |  |  |  | 0.56** | 0.03 |
| Total | -0.06* | 0.03 | -0.05 | 0.03 | -0.06** | 0.02 |
| Externalizing (Baseline) |  |  |  |  | 0.62** | 0.03 |
| **WJ-III** |  |  |  |  |  |  |
| Applied problems | -0.05* | 0.02 | -0.03 | 0.02 | -0.04** | 0.01 |
| Externalizing (Baseline) |  |  |  |  | 0.62** | 0.03 |
| Letter Identification | -0.06** | 0.02 | -0.05 | 0.02 | -0.03** | 0.01 |
| Externalizing (Baseline) |  |  |  |  | 0.61** | 0.03 |
| Passage Comp.<sup>+</sup> | -0.07 | 0.03 | -0.05 | 0.03 | -0.01 | 0.02 |
| Externalizing (Baseline) |  |  |  |  | 0.58** | 0.04 |
| Total score | -0.07** | 0.02 | -0.05 | 0.02 | -0.04** | 0.01 |
| Externalizing (Baseline) |  |  |  |  | 0.62** | 0.03 |
| **Sum score** |  |  |  |  |  |  |
| Composite Intelligence | -0.08** | 0.02 | -0.06* | 0.02 | -0.05** | 0.01 |
| Externalizing (Baseline) |  |  |  |  | 0.62** | 0.03 |

\*p<.05, \*\*p<.01
Note. Model 1 regresses outcome on intelligence at baseline, Model 2 adds covariates, and Model 3 regresses outcome at Time 2 onto outcome at Time 1, IQ index, and covariates. <sup>+</sup>Models with the Passage Comprehension subtest have a sample size of n = 1566 because the subtest was not administered to participants age 12+.

**Table 4:** Regression models predicting internalizing CBCL conditions at time 2

|  | Model 1 |  | Model 2 |  | Model 3 |  |
| --- | --- | --- | --- | --- | --- | --- |
|  | *b* | *SE* | *b* | *SE* | *b* | *SE* |
| **K-BIT** |  |  |  |  |  |  |
| Matrices | -0.02 | 0.02 | -0.01 | 0.02 | -0.04* | 0.02 |
| Internalizing (Baseline) |  |  |  |  | 0.61** | 0.03 |
| Verbal | -0.02 | 0.02 | -0.02 | 0.02 | -0.04* | 0.01 |
| Internalizing (Baseline) |  |  |  |  | 0.60 | 0.03 |
| Total | -0.02 | 0.02 | -0.02 | 0.02 | -0.05* | 0.02 |
| Internalizing (Baseline) |  |  |  |  | 0.60** | 0.03 |
| **WJ-III** |  |  |  |  |  |  |
| Applied problems | -0.03 | 0.02 | -0.01 | 0.02 | -0.02 | 0.01 |
| Internalizing (Baseline) |  |  |  |  | 0.59 | 0.03 |
| Letter Identification | -0.04* | 0.02 | -0.02 | 0.02 | -0.03* | 0.01 |
| Internalizing (Baseline) |  |  |  |  | 0.59** | 0.03 |
| Passage Comp.<sup>+</sup> | -0.06 | 0.03 | -0.04 | 0.04 | -0.02 | 0.02 |
| Internalizing (Baseline) |  |  |  |  | 0.54** | 0.04 |
| Total | -0.04* | 0.02 | -0.02 | 0.02 | -0.04* | 0.01 |
| Internalizing (Baseline) |  |  |  |  | 0.59** | 0.03 |
| **Sum score** |  |  |  |  |  |  |
| Composite Intelligence | -0.04 | 0.02 | -0.03 | 0.02 | -0.05* | 0.01 |
| Internalizing (Baseline) |  |  |  |  | 0.60 | 0.03 |

\*p<.05, \*\*p<.01
Note. Model 1 regresses outcome on intelligence at baseline, Model 2 adds covariates, and Model 3 regresses outcome at Time 2 onto outcome at Time 1, IQ indice, and covariates. <sup>+</sup>Models with the Passage Comprehension subtest have a sample size of n = 1566 because the subtest was not administered to participants age 12+.

**Table 5:** Regression models predicting total CBCL behavioral conditions at time 2

|  | Model 1 |  | Model 2 |  | Model 3 |  |
| --- | --- | --- | --- | --- | --- | --- |
|  | *b* | *SE* | *b* | *SE* | *b* | *SE* |
| **K-BIT** |  |  |  |  |  |  |
| Matrices | -0.05 | 0.03 | -0.04 | 0.03 | -0.06** | 0.02 |
| Total Behavior (Baseline) |  |  |  |  | 0.67** | 0.03 |
| Verbal | -0.05 | 0.03 | -0.05 | 0.03 | -0.06** | 0.02 |
| Total Behavior (Baseline) |  |  |  |  | 0.66** | 0.03 |
| Total | -0.06 | 0.03 | -0.06 | 0.04 | -0.08** | 0.02 |
| Total Behavior (Baseline) |  |  |  |  | 0.67** | 0.03 |
| **WJ-III** |  |  |  |  |  |  |
| Applied problems | -0.06* | 0.02 | -0.04 | 0.03 | -0.04** | 0.01 |
| Total Behavior (Baseline) |  |  |  |  | 0.66** | 0.04 |
| Letter Identification | -0.07** | 0.02 | -0.05 | 0.02 | -0.04** | 0.01 |
| Total Behavior (Baseline) |  |  |  |  | 0.66** | 0.04 |
| Passage Comp.<sup>+</sup> | -0.09 | 0.04 | -0.06 | 0.05 | -0.02 | 0.02 |
| Total Behavior (Baseline) |  |  |  |  | 0.62** | 0.04 |
| Total | -0.08** | 0.03 | -0.06 | 0.03 | -0.05** | 0.01 |
| Total Behavior (Baseline) |  |  |  |  | 0.66 | 0.04 |
| **Sum Score** |  |  |  |  |  |  |
| Composite Intelligence | -0.09** | 0.03 | -0.07* | 0.03 | -0.07** | 0.02 |
| Total Behavior (Baseline) |  |  |  |  | 0.66** | 0.03 |

\*p<.05, \*\*p<.01
Note. Model 1 regresses outcome on intelligence at baseline, Model 2 adds covariates, and Model 3 regresses outcome at Time 2 onto outcome at Time 1, IQ indice, and covariates. <sup>+</sup>Models with the Passage Comprehension subtest have a sample size of n = 1566 because the subtest was not administered to participants age 12+.

### Do associations between intelligence and psychopathology hold after controlling for important covariates?

Model 2 across Tables 3-5 presents the results of regression analyses predicting the relevant psychopathology variable one year later while controlling for child age, race, and sex, as well as an indicator of poverty^2^. Virtually all associations found between intelligence and psychopathology in the various Model 1’s were rendered nonsignificant after the inclusion of these covariates. The only associations which remained, were inverse associations between our composite intelligence variable and externalizing and total behavior problem ratings.

### Does intelligence predict changes in psychopathology after controlling for potential confounds?

Model 3 in our regression tables (Tables 3-5) presents the results of regression analyses predicting psychopathology one year later, while controlling for our demographic covariates *and* baseline levels of psychopathology, allowing for an examination of the association between intelligence and changes in psychopathology over time. All intelligence variables – except WJ-III Passage Comprehension – were significantly, negatively related to externalizing symptoms and total behavior problems at time 2 after controlling for baseline externalizing. Similarly, all indicators of intelligence except for WJ-III Applied Problems and Passage Comprehension were negatively related to changes in internalizing symptoms. Overall, the results obtained in these models suggest that lower levels of intelligence are consistently associated with increases in psychopathology over time – at least over the course of one year.

Finally, and owing to possible concerns about spuriousness, some sensitivity analysis regarding a determination of substantiation (of abuse/neglect) and placement following the investigation was conducted by entering these variables into the analyses as additional covariates^3^. In almost every model, the introduction of these variables did not substantively alter the relationship between intelligence and behavior. However, the relationship between intelligence as measured by the KBIT matrices score and internalizing behavioral problems (top row of Table 4) was rendered insignificant. As one last approach to examining the effects of disadvantage on the association between intelligence and psychopathology, we estimated regression models testing for an interaction between intelligence and poverty^4^. These analyses did not substantively change our findings.

## Discussion

The association between internalizing and externalizing problems has been well known to, and widely discussed by psychopathology researchers for some time (Caspi et al., 2014; Del Giudice, 2016). Increasingly, scholars have attempted to understand the source of the comorbidity in order to better gauge whether certain risk factors are common across both types of disorders. At the same time, intelligence researchers have reported well-replicated associations between indicators of general intelligence and various deleterious outcomes including overt antisocial behavior, violence, aggression, and low impulse control (see citations above). In other words, there exists evidence for a link between intelligence and externalizing behaviors but less so for internalizing problems. Our study was intended to examine internalizing alongside externalizing, as well as test if the relationship between intelligence and psychopathology held in a highly at-risk and disadvantaged sample of American participants.

Intelligence, as measured by both the K-BIT and WJ-III, in the present study of at-risk youth seemed to be more consistently associated with externalizing and total behavioral problems rather than internalizing problems. Interestingly, the K-BIT did not seem to predict externalizing problems as robustly as the WJ-III composite, which it should be noted, generally (though not entirely) assesses variation in achievement and crystalized intelligence (see Cattell, 1967; Ritchie, 2015 for details on the distinction between intelligence types). A less consistent association emerged for internalizing problems and the indicators of intelligence, though the WJ-III did evince sporadic evidence of correlation with the internalizing items of the CBCL. Finally, the total CBCL was associated with lower scores on the composite intelligence measure in the sample. Thus, the results did deviate from expectation in some instances. It does seem worth noting, though, that some general picture emerged of an association between intelligence and externalizing problems, along with a much less consistent relationship for internalizing problems, and an overall association for behavioral problems (broadly defined) and intelligence. The smaller effect of intelligence on internalizing compared to externalizing behaviors, it should also be mentioned, is consistent with previous associations found in more advantaged populations (Flouri, Midouhas, & Joshi, 2015).

Findings pertaining to the WJ-III warrant some additional comment, as they may suggest a role for crystalized intelligence in general, and reading performance in particular, in the origins of externalizing problem behavior in at-risk children. This observation is in line with prior research finding an association between verbal ability and delinquency, in part because increased verbal ability might permit greater frustration tolerance, and solving interpersonal conflict via communication as opposed to violent outbursts (Bellair, McNulty, and Piquero, 2016; Moffitt, Lynam, and Silva, 1994). From a more pragmatic perspective, this finding (very tentatively) suggests the possibility that reading intervention for struggling readers in this at-risk population *may* represent one possible avenue for reducing externalizing behavior (Vacca, 2008). Such a recommendation should be tempered, however, as the results of the K-BIT verbal test (as well as the WJ-III in various models) were occasionally at odds with this, generally only emerging as significant in models examining changes in behavior over time. Any concrete policy recommendations at this juncture would be decidedly premature.

Prior to concluding, there are a number of limitations and considerations that are important to mention. First, although we primarily consider the composition of our sample (i.e., an at-risk portion of American children) a strength, our findings cannot be assumed to generalize to the broader population. Owing to the relatively little effort aimed at examining the phenotypic association between indicators of intelligence and behavioral outcomes in such populations, though, we view the characteristics of the current sample as a desirable quality. Second, the assessment of externalizing and internalizing symptoms is based solely on caregiver reports and is therefore subject to the usual threats of social desirability bias and over and under reporting. Future research would benefit from the use of multi-method assessment of psychopathology. Another consideration involves the tendency for the statistically significant association of intelligence and the outcome variables to dissipate entirely in Model 2 when other key control variables were introduced. While this may be in part due to small increases in variation attributable to these covariates, the actual effect size was not substantially attenuated across models.

Moreover, and as is often the case in associational research, issues such as collider bias and residual confounding cannot be ruled out. Collider bias, in particular, refers to situations in which certain covariates included in a multivariate model are *directly impacted* by both the key independent variable and the dependent variable (Rohrer, 2018). When this is the case, it can bias the influence of the focal independent variable in the study. In our case, we included covariates for age, sex, race, and (family) poverty. Given that our focus was on measurement collected in children, collider bias seems less likely given the limited influence of child traits on covariates like parental SES in particular (had our sample been adults, this may have presented a more serious problem). Nonetheless, specifying a potentially causal model in associational research requires careful exploration and cross-validation in the form of replication using designs that permit stronger causal inference than what we can offer. Additionally, the use of tools such as directed acyclic graphs (DAGs) to better specify potential casual pathways, and to block back door effects, would also be useful for this topic (see Rohrer, 2018 for a general outline on these topics).

Finally, the primary independent and dependent variables in the study are moderately to highly heritable constructs (Plomin & Deary, 2015), however, the nature of our data did not permit us to model the effects of heritability directly. This is an important issue, because as prior research has noted (Barnes, Boutwell, Gibson, Beaver & Wright, 2014), even a moderate genetic correlation left unaccounted for may fully confound a phenotypic correlation. Thus, it remains to be seen whether the phenotypic association between intelligence and externalizing/internalizing behaviors observed herein can withstand correction for genetic confounding (for review of genetic influences on intelligence and psychopathology, see Smoller et al., 2018).

In conclusion, we observed a set of findings in an at-risk US sample that seem—at times—in line with the limited prior research on this topic, though there were some divergences too. Higher intelligence was, in various models, associated with lower instances of child externalizing behavioral problems. This same relationship was also observed with internalizing behaviors – including feelings of anxiety and depression - but in substantially fewer of the models. Ultimately, our paper offers additional insight into the association (and perhaps lack thereof, in some cases) between indicators of intelligence and problem behaviors. To the extent that our findings replicate in other samples, using other measures of intelligence and psychopathological outcomes remains an open and important question for future researchers to address.

## Footnotes

1 As in the current study, Harpur et al. (2015) included both substantiated and unsubstantiated cases of child abuse and neglect.

2 Tables presenting all effect sizes and statistics for the full models, including covariates, are presented in the supplementary file.

3 We would like to thank our anonymous reviewers for mentioning this point and suggesting the additional analyses.

4 Due to 25% of the sample being in foster care at the time of data collection, these participants were removed as we were unable to assess their exposure to poverty prior to foster care placement. As a result, we stratified the remaining participants into two categories – in poverty vs not in poverty – to conserve statistical power.

