## Supplemental Tables for "The Association of Externalizing and Internalizing Problems with Indicators of Intelligence in a Sample of At-Risk Children"

S2: Regression models predicting externalizing (CBCL) conditions at Wave 2

| | Model 1 | | Model 2 | | Model 3 | |
|---|---|---|---|---|---|---|
| | *b* | *SE* | *b* | *SE* | *b* | *SE* |
| **K-BIT** | | | | | | |
| Matrices | -0.06* | 0.03 | -0.04 | 0.03 | -0.06** | 0.02 |
| Age (y) | | | 0.40** | 0.13 | 0.01 | 0.09 |
| Female | | | -1.87* | 0.78 | -0.42 | 0.71 |
| Race | | | | | | |
| Non-Hispanic White | | | | | | |
| Non-Hispanic Black | | | -1.78 | 1.19 | -1.03 | 0.93 |
| Hispanic | | | -3.78** | 0.92 | -2.67** | 0.80 |
| Other | | | -0.52 | 1.29 | -1.05 | 0.93 |
| Family poverty | | | | | | |
| <50% | | | 0.52 | 1.20 | 0.79 | 0.62 |
| 51%-100% | | | -0.37** | 1.02 | -0.24** | 0.86 |
| 101%-200% | | | | | | |
| >200% | | | -2.83* | 1.15 | -0.97** | 1.21 |
| Externalizing (Baseline) | | | | | 0.62** | 0.03 |
| **K-BIT** | | | | | | |
| Verbal | -0.04 | 0.02 | -0.04 | 0.03 | -0.04* | 0.02 |
| Age (y) | | | 0.40** | 0.13 | 0.03* | 0.09 |
| Female | | | -1.87* | 0.75 | -0.4 | 0.70 |
| Race | | | | | | |
| Non-Hispanic White | | | | | | |
| Non-Hispanic Black | | | -2.07 | 1.22 | -1.41 | 0.98 |
| Hispanic | | | -4.04** | .97 | -3.16** | 0.83 |
| Other | | | -0.94 | 1.27 | -1.41 | 0.92 |
| Family poverty | | | | | | |
| <50% | | | 0.52 | 1.12 | 0.86 | 0.59 |
| 51%-100% | | | -0.53 | 0.98 | -0.34 | 0.86 |
| 101%-200% | | | | | | |

| | | | | | | |
|---|---|---|---|---|---|---|
| >200% | | | -2.73** | 1.09 | -0.93 | 1.16 |
| Externalizing (Baseline) | | | | | 0.56** | 0.03 |
| **K-BIT** | | | | | | |
| Total | -0.06* | 0.03 | -0.05 | 0.03 | -0.06** | 0.02 |
| Age (y) | | | 0.39** | 0.13 | 0.01 | 0.09 |
| Female | | | -1.86* | 0.78 | -0.41 | 0.73 |
| Race | | | | | | |
| Non-Hispanic White | | | | | | |
| Non-Hispanic Black | | | -1.93 | 1.18 | -1.25 | 0.95 |
| Hispanic | | | -4.0** | 0.95 | -2.97** | 0.82 |
| Other | | | -0.55 | 1.28 | -1.13 | 0.94 |
| Family poverty | | | | | | |
| <50% | | | 0.38 | 1.19 | 0.71 | 0.63 |
| 51%-100% | | | -0.54 | 1.03 | -0.31 | 0.88 |
| 101%-200% | | | | | | |
| >200% | | | -2.90* | 1.16 | -0.97 | 1.22 |
| Externalizing (Baseline) | | | | | 0.62** | 0.03 |
| **WJ-III** | | | | | | |
| Applied problems | -0.05* | 0.02 | -0.03 | 0.02 | -0.04** | 0.01 |
| Age (y) | | | 0.28* | 0.13 | -0.04 | 0.09 |
| Female | | | -2.16** | 0.79 | -0.43 | 0.76 |
| Race | | | | | | |
| Non-Hispanic White | | | | | | |
| Non-Hispanic Black | | | -2.30 | 1.29 | -1.24 | 0.98 |
| Hispanic | | | -4.44** | 0.91 | -3.04** | 0.79 |
| Other | | | -0.52 | 1.25 | -1.12 | 0.92 |
| Family poverty | | | | | | |
| <50% | | | 1.03 | 0.95 | 1.34* | 0.57 |
| 51%-100% | | | -0.44 | 1.06 | 0.06 | 0.89 |
| 101%-200% | | | | | | |
| >200% | | | -3.26** | 1.18 | -1.00 | 1.11 |

| | | | | | | |
|---|---|---|---|---|---|---|
| Externalizing (Baseline) | | | | | 0.62** | 0.03 |
| **WJ-III** | | | | | | |
| Letter Identification | -0.06** | 0.02 | -0.05 | 0.02 | -0.03** | 0.01 |
| Age (y) | | | 0.28* | 0.13 | -0.01 | 0.09 |
| Female | | | -2.10** | 0.79 | -0.38 | 0.77 |
| Race | | | | | | |
| Non-Hispanic White | | | | | | |
| Non-Hispanic Black | | | -2.22 | 1.27 | -1.23 | 0.99 |
| Hispanic | | | -4.29** | 0.94 | -3.05** | 0.78 |
| Other | | | -0.25 | 1.31 | -1.09 | 0.97 |
| Family poverty | | | | | | |
| <50% | | | 0.98 | 0.95 | 1.39* | 0.57 |
| 51%-100% | | | -0.33 | 1.06 | 0.13 | 0.9 |
| 101%-200% | | | | | | |
| >200% | | | -3.11** | 1.14 | -1.0 | 1.12 |
| Externalizing (Baseline) | | | | | 0.61** | 0.03 |
| **WJ-III** | | | | | | |
| Passage Comp.+ | -0.07 | 0.03 | -0.05 | 0.03 | -0.01 | 0.02 |
| Age (y) | | | -0.07 | 0.25 | -0.31 | 0.20 |
| Female | | | -2.59* | 1.05 | -0.07 | 0.98 |
| Race | | | | | | |
| Non-Hispanic White | | | | | | |
| Non-Hispanic Black | | | -3.28 | 1.77 | -2.67* | 1.32 |
| Hispanic | | | -2.81* | 1.16 | -1.79 | 1.10 |
| Other | | | 0.12 | 2.83 | -1.04 | 1.96 |
| Family poverty | | | | | | |
| <50% | | | 1.83 | 1.5 | 1.56 | 0.92 |
| 51%-100% | | | 1.77 | 1.08 | 1.27 | 0.88 |
| 101%-200% | | | | | | |
| >200% | | | -3.17 | 1.66 | -1.87 | 1.41 |
| Externalizing (Baseline) | | | | | 0.58** | 0.04 |

| | | | | | | |
|---|---|---|---|---|---|---|
| **WJ-III** | | | | | | |
| Total score | -0.07** | 0.02 | -0.05 | 0.02 | -0.04** | 0.01 |
| Age (y) | | | 0.27* | 0.13 | -0.03 | 0.10 |
| Female | | | -2.11** | 0.78 | -0.36 | 0.75 |
| Race | | | | | | |
| Non-Hispanic White | | | | | | |
| Non-Hispanic Black | | | -2.23 | 1.26 | -1.25 | 0.97 |
| Hispanic | | | -4.30** | 0.92 | -3.04** | 0.77 |
| Other | | | -0.30 | 1.28 | -1.04 | 0.94 |
| Family poverty | | | | | | |
| <50% | | | 0.98 | 0.95 | 1.36* | 0.57 |
| 51%-100% | | | -0.35 | 1.06 | 0.08 | 0.88 |
| 101%-200% | | | | | | |
| >200% | | | -3.15** | 1.17 | -0.98 | 1.12 |
| Externalizing (Baseline) | | | | | 0.62** | 0.03 |
| **Sum Score** | | | | | | |
| Composite Intelligence | -0.08** | 0.02 | -0.06* | 0.02 | -0.05** | 0.01 |
| Age (y) | | | 0.37** | 0.13 | 0.01 | 0.09 |
| Female | | | -1.86* | 0.73 | -0.37 | 0.70 |
| Race | | | | | | |
| Non-Hispanic White | | | | | | |
| Non-Hispanic Black | | | -1.91 | 1.18 | -1.17 | 0.91 |
| Hispanic | | | -3.83** | 0.89 | -2.88** | 0.8 |
| Other | | | -0.65** | 1.28 | -1.13 | 0.90 |
| Family poverty | | | | | | |
| <50% | | | 0.30 | 1.11 | 0.67 | 0.55 |
| 51%-100% | | | -0.53** | 0.96 | -0.36 | 0.83 |
| 101%-200% | | | | | | |
| >200% | | | -2.70** | 1.05 | -0.92 | 1.16 |
| Externalizing (Baseline) | | | | | 0.62** | 0.03 |

*Note*. Model 1 regresses outcome on intelligence at baseline, Model 2 adds covariates, and and Model 3 regresses outcome at Time 2 onto outcome at Time 1, IQ indice, and covariates. + Models with the Passage Comprehension subtest have a sample size of n = 1566 because the subtest was not administered to participants age 12+.

S3: Regression models predicting internalizing (CBCL) conditions at Wave 2

| | Model 1 | | Model 2 | | Model 3 | |
|---|---|---|---|---|---|---|
| | *b* | *SE* | *B* | *SE* | *b* | *SE* |
| **K-BIT** | | | | | | |
| Matrices | -0.02 | 0.02 | -0.01 | 0.02 | -0.04* | 0.02 |
| Age (y) | | | 0.33* | 0.12 | 0.09 | 0.08 |
| Female | | | -1.40 | 0.80 | -0.33 | 0.65 |
| Race | | | | | | |
| Non-Hispanic White | | | | | | |
| Non-Hispanic Black | | | -3.65** | 1.03 | -2.40** | 0.80 |
| Hispanic | | | -1.85 | 1.15 | -2.32** | 0.84 |
| Other | | | 0.44 | 1.30 | -1.34 | 1.03 |
| Family poverty | | | | | | |
| <50% | | | 0.63 | 1.43 | 0.67 | 0.78 |
| 51%-100% | | | 0.17 | 1.10 | 0.28 | 0.86 |
| 101%-200% | | | | | | |
| >200% | | | -2.47* | 1.15 | -0.32** | 1.04 |
| Internalizing (Baseline) | | | | | 0.61** | 0.03 |
| **K-BIT** | | | | | | |
| Verbal | -0.02 | 0.02 | -0.02 | 0.02 | -0.04* | 0.01 |
| Age (y) | | | 0.31 | 0.12 | 0.08 | 0.08 |
| Female | | | -1.42 | 0.76 | -0.32 | 0.64 |
| Race | | | | | | |
| Non-Hispanic White | | | | | | |
| Non-Hispanic Black | | | -3.55** | 1.05 | -2.55** | 0.79 |
| Hispanic | | | -1.90 | 1.14 | -2.68** | 0.78 |
| Other | | | 0.28 | 1.24 | -1.55 | 0.99 |
| Family poverty | | | | | | |
| <50% | | | 0.70 | 1.37 | 0.74 | 0.76 |

| | | | | | | |
|---|---|---|---|---|---|---|
| 51%-100% | | | 0.34 | 1.10 | 0.37 | 0.86 |
| 101%-200% | | | | | | |
| >200% | | | -2.16 | 1.10 | -0.01** | 0.98 |
| Internalizing (Baseline) | | | | | 0.60 | 0.03 |
| **K-BIT** | | | | | | |
| Total | -0.02 | 0.02 | -0.02 | 0.02 | -0.05* | 0.02 |
| Age (y) | | | 0.32* | 0.13 | 0.07 | 0.08 |
| Female | | | -1.35 | 0.80 | -0.33 | 0.66 |
| Race | | | | | | |
| Non-Hispanic White | | | | | | |
| Non-Hispanic Black | | | -3.66** | 1.04 | -2.53** | 0.80 |
| Hispanic | | | -1.88** | 1.17 | -2.50** | 0.82 |
| Other | | | 0.51 | 1.28 | -1.35 | 1.02 |
| Family poverty | | | | | | |
| <50% | | | 0.58 | 1.43 | 0.59 | 0.78 |
| 51%-100% | | | 0.14 | 1.12 | 0.24 | 0.87 |
| 101%-200% | | | | | | |
| >200% | | | -2.44* | 1.16 | -0.27 | 1.04 |
| Internalizing (Baseline) | | | | | 0.60** | 0.03 |
| **WJ-III** | | | | | | |
| Applied problems | -0.03 | 0.02 | -0.01 | 0.02 | -0.02 | 0.01 |
| Age (y) | | | 0.45** | 0.13 | 0.22* | 0.09 |
| Female | | | -1.91* | 0.89 | -0.45 | 0.78 |
| Race | | | | | | |
| Non-Hispanic White | | | | | | |
| Non-Hispanic Black | | | -3.75** | 1.13 | -2.33** | 0.82 |
| Hispanic | | | -2.30* | 1.11 | -2.51** | 0.89 |
| Other | | | 0.33 | 1.32 | -1.42 | 1.01 |
| Family poverty | | | | | | |
| <50% | | | 1.12 | 1.16 | 1.03 | 0.80 |
| 51%-100% | | | 0.28 | 1.10 | 0.45 | 0.80 |
| 101%-200% | | | | | | |
| >200% | | | -2.37* | 1.06 | 0.15 | 0.96 |

| | | | | | | |
|---|---|---|---|---|---|---|
| Internalizing (Baseline) | | | | | 0.59 | 0.03 |
| **WJ-III** | | | | | | |
| Letter Identification | -0.04* | 0.02 | -0.02 | 0.02 | -0.03* | 0.01 |
| Age (y) | | | 0.43** | 0.13 | 0.22** | 0.08 |
| Female | | | -1.92* | 0.90 | -0.38 | 0.80 |
| Race | | | | | | |
| Non-Hispanic White | | | | | | |
| Non-Hispanic Black | | | -3.68** | 1.11 | -2.25** | 0.82 |
| Hispanic | | | -2.13 | 1.13 | -2.41** | 0.90 |
| Other | | | 0.53 | 1.3 | -1.24 | 1.00 |
| Family poverty | | | | | | |
| <50% | | | 1.06 | 1.17 | 1.01 | 0.78 |
| 51%-100% | | | 0.40 | 1.10 | 0.51 | 0.76 |
| 101%-200% | | | | | | |
| >200% | | | -2.25* | 1.09 | 0.22 | 0.96 |
| Internalizing (Baseline) | | | | | 0.59** | 0.03 |
| **WJ-III** | | | | | | |
| Passage Comp.+ | -0.06 | 0.03 | -0.04 | 0.04 | -0.02 | 0.02 |
| Age (y) | | | 0.38 | 0.25 | 0.19 | 0.17 |
| Female | | | -3.36** | 1.15 | -1.05 | 1.09 |
| Race | | | | | | |
| Non-Hispanic White | | | | | | |
| Non-Hispanic Black | | | -4.12* | 1.57 | -2.90* | 1.08 |
| Hispanic | | | -1.40 | 1.29 | -2.11* | 1.04 |
| Other | | | 1.21 | 2.25 | -0.04 | 1.70 |
| Family poverty | | | | | | |
| <50% | | | 2.51 | 1.64 | 2.07 | 1.15 |
| 51%-100% | | | 1.88 | 1.13 | 1.62 | 0.99 |
| 101%-200% | | | | | | |
| >200% | | | -0.43 | 1.25 | 1.32 | 0.96 |
| Internalizing (Baseline) | | | | | 0.54** | 0.04 |
| **WJ-III** | | | | | | |
| Total | -0.04* | 0.02 | -0.02 | 0.02 | -0.04* | 0.01 |

| | | | | | | |
|---|---|---|---|---|---|---|
| Age (y) | | | 0.43** | 0.13 | 0.21* | 0.08 |
| Female | | | -1.92* | 0.89 | -0.40 | 0.78 |
| Race | | | | | | |
| Non-Hispanic White | | | | | | |
| Non-Hispanic Black | | | -3.66** | 1.11 | -2.26** | 0.81 |
| Hispanic | | | -2.12 | 1.13 | -2.42** | 0.89 |
| Other | | | 0.52 | 1.31 | -1.23 | 1.01 |
| Family poverty | | | | | | |
| <50% | | | 1.04 | 1.17 | 0.98 | 0.78 |
| 51%-100% | | | 0.40 | 1.11 | 0.49 | 0.77 |
| 101%-200% | | | | | | |
| >200% | | | -2.26* | 1.08 | 0.26 | 0.96 |
| Internalizing (Baseline) | | | | | 0.59** | 0.03 |
| **Sum Score** | | | | | | |
| Composite Intelligence | -0.04 | 0.02 | -0.03 | 0.02 | -0.05* | 0.01 |
| Age (y) | | | 0.30* | 0.13 | 0.07 | 0.08 |
| Female | | | -1.33 | 0.77 | -0.20 | 0.66 |
| Race | | | | | | |
| Non-Hispanic White | | | | | | |
| Non-Hispanic Black | | | -3.42** | 1.04 | -2.25** | 0.78 |
| Hispanic | | | -1.66 | 1.13 | -2.32** | 0.80 |
| Other | | | 0.53 | 1.28 | -1.19 | 1.00 |
| Family poverty | | | | | | |
| <50% | | | 0.29 | 1.38 | 0.41 | 0.79 |
| 51%-100% | | | 0.35 | 1.07 | 0.40 | 0.84 |
| 101%-200% | | | | | | |
| >200% | | | -2.25* | 1.08 | -0.03 | 0.98 |
| Internalizing (Baseline) | | | | | 0.60** | 0.03 |

*p<.05, **p<.01

*Note*. Model 1 regresses outcome on intelligence at baseline, Model 2 adds covariates, and and Model 3 regresses outcome at Time 2 onto outcome at Time 1, IQ indice, and covariates. + Models with the Passage Comprehension subtest have a sample size of n = 1566 because the subtest was not administered to participants age 12+.

S4: Regression models predicting total (CBCL) behavioral conditions at Wave 2

| | Model 1 | | Model 2 | | Model 3 | |
|---|---|---|---|---|---|---|
| | *b* | *SE* | *b* | *SE* | *b* | *SE* |
| **K-BIT** | | | | | | |
| Matrices | -0.05 | 0.02 | -0.04 | 0.03 | -0.06** | 0.02 |
| Age (y) | | | 0.39** | 0.14 | 0.01 | 0.08 |
| Female | | | -2.11* | 0.86 | -0.62 | 0.70 |
| Race | | | | | | |
| Non-Hispanic White | | | | | | |
| Non-Hispanic Black | | | -3.17* | 1.25 | -1.93* | 0.91 |
| Hispanic | | | -3.48* | 1.03 | -2.67** | 0.77 |
| Other | | | -0.51 | 1.48 | -1.81 | 1.12 |
| Family poverty | | | | | | |
| <50% | | | 0.32 | 1.4 | 0.45 | 0.63 |
| 51%-100% | | | -0.06 | 1.09 | -0.15 | 0.85 |
| 101%-200% | | | | | | |
| >200% | | | -2.71* | 1.12 | -0.59 | 1.07 |
| Total behavior (Baseline) | | | | | 0.67** | 0.03 |
| **K-BIT** | | | | | | |
| Verbal | -0.05 | 0.02 | -0.05 | 0.03 | -0.06** | 0.02 |
| Age (y) | | | 0.38* | 0.15 | 0.01 | 0.09 |
| Female | | | -2.22** | 0.83 | -0.67 | 0.70 |
| Race | | | | | | |
| Non-Hispanic White | | | | | | |
| Non-Hispanic Black | | | -3.41** | 1.30 | -2.33* | 0.91 |
| Hispanic | | | -3.74** | 1.06 | -3.24** | 0.76 |
| Other | | | -0.92 | 1.43 | -2.15 | 1.10 |
| Family poverty | | | | | | |
| <50% | | | 0.27 | 1.32 | 0.52 | 0.60 |
| 51%-100% | | | -0.09 | 1.07 | -0.18 | 0.84 |

| | | | | | | |
|---|---|---|---|---|---|---|
| 101%-200% | | | | | | |
| >200% | | | -2.50* | 1.05 | -0.41 | 1.01 |
| Total behavior (Baseline) | | | | | 0.66** | 0.03 |
| **K-BIT** | | | | | | |
| Total | -0.06 | 0.03 | -0.06 | 0.04 | -0.08** | 0.02 |
| Age (y) | | | 0.38* | 0.15 | -0.01 | 0.09 |
| Female | | | -2.11* | 0.87 | -0.65 | 0.72 |
| Race | | | | | | |
| Non-Hispanic White | | | | | | |
| Non-Hispanic Black | | | -3.35** | 1.26 | -2.21* | 0.91 |
| Hispanic | | | -3.69** | 1.06 | -2.98** | 0.77 |
| Other | | | -0.48 | 1.46 | -1.86 | 1.13 |
| Family poverty | | | | | | |
| <50% | | | 0.20 | 1.40 | 0.37 | 0.65 |
| 51%-100% | | | -0.18 | 1.10 | -0.20 | 0.86 |
| 101%-200% | | | | | | |
| >200% | | | -2.69* | 1.14 | -0.49 | 1.07 |
| Total behavior (Baseline) | | | | | 0.67** | 0.03 |
| **WJ-III** | | | | | | |
| Applied problems | -0.06* | 0.02 | -0.04 | 0.03 | -0.04** | 0.01 |
| Age (y) | | | 0.33* | 0.16 | 0.01 | 0.10 |
| Female | | | -2.65** | 0.86 | -0.74 | 0.79 |
| Race | | | | | | |
| Non-Hispanic White | | | | | | |
| Non-Hispanic Black | | | -3.54* | 1.41 | -1.94* | 0.94 |
| Hispanic | | | -4.07** | 0.97 | -2.98** | 0.73 |
| Other | | | -0.69 | 1.56 | -1.95 | 1.16 |
| Family poverty | | | | | | |
| <50% | | | 0.67 | 1.10 | 0.88 | 0.57 |
| 51%-100% | | | 0.01** | 1.16 | 0.19 | 0.82 |

| | | | | | | |
|---|---|---|---|---|---|---|
| 101%-200% | | | | | | |
| >200% | | | -2.98** | 1.12 | -0.34 | 0.97 |
| Total behavior (Baseline) | | | | | 0.66** | 0.04 |
| **WJ-III** | | | | | | |
| Letter Identification | -0.07** | 0.02 | -0.05 | 0.02 | -0.04** | 0.01 |
| Age (y) | | | 0.33* | 0.16 | 0.03 | 0.10 |
| Female | | | -2.55** | 0.84 | -0.65 | 0.79 |
| Race | | | | | | |
| Non-Hispanic White | | | | | | |
| Non-Hispanic Black | | | -3.47* | 1.38 | -1.91* | 0.93 |
| Hispanic | | | -4.00** | 0.96 | -3.00** | 0.73 |
| Other | | | -0.44 | 1.60 | -1.87 | 1.22 |
| Family poverty | | | | | | |
| <50% | | | 0.63 | 1.10 | 0.93 | 0.57 |
| 51%-100% | | | 0.05* | 1.15 | 0.23 | 0.82 |
| 101%-200% | | | | | | |
| >200% | | | -2.76* | 1.11 | -0.27 | 1.0 |
| Total behavior (Baseline) | | | | | 0.66** | 0.04 |
| **WJ-III** | | | | | | |
| Passage Comp.+ | -0.09 | 0.04 | -0.06 | 0.05 | -0.02 | 0.02 |
| Age (y) | | | 0.07 | 0.27 | -0.15 | 0.20 |
| Female | | | -3.51** | 1.15 | -0.78 | 1.05 |
| Race | | | | | | |
| Non-Hispanic White | | | | | | |
| Non-Hispanic Black | | | -4.37* | 2.01 | -3.27* | 1.30 |
| Hispanic | | | -2.80* | 1.09 | -2.13* | 1.07 |
| Other | | | 0.62 | 3.18 | -1.11 | 2.29 |
| Family poverty | | | | | | |
| <50% | | | 2.32 | 1.74 | 1.96* | 0.90 |
| 51%-100% | | | 2.61* | 1.17 | 1.86* | 0.89 |

| | | | | | | |
|---|---|---|---|---|---|---|
| 101%-200% | | | | | | |
| >200% | | | -1.60 | 1.66 | 0.08 | 1.21 |
| Total behavior (Baseline) | | | | | 0.62** | 0.04 |
| **WJ-III** | | | | | | |
| Total | -0.08** | 0.03 | -0.06 | 0.03 | -0.05** | 0.01 |
| Age (y) | | | 0.32* | 0.16 | 0.02 | 0.10 |
| Female | | | -2.55** | 0.84 | -0.63 | 0.77 |
| Race | | | | | | |
| Non-Hispanic White | | | | | | |
| Non-Hispanic Black | | | -3.44** | 1.38 | -1.92* | 0.92 |
| Hispanic | | | -3.98** | 0.96 | -3.00** | 0.71 |
| Other | | | -0.43 | 1.58 | -1.81 | 1.20 |
| Family poverty | | | | | | |
| <50% | | | 0.60 | 1.10 | 0.88 | 0.57 |
| 51%-100% | | | 0.05 | 1.15 | 0.18 | 0.81 |
| 101%-200% | | | | | | |
| >200% | | | -2.83* | 1.12 | -0.27 | 0.98 |
| Total behavior (Baseline) | | | | | 0.66 | 0.04 |
| **Sum Score** | | | | | | |
| Composite Intelligence | -0.09** | 0.03 | -0.07* | 0.03 | -0.07** | 0.02 |
| Age (y) | | | 0.35* | 0.15 | -0.01 | 0.09 |
| Female | | | -2.11* | 0.81 | -0.56 | 0.69 |
| Race | | | | | | |
| Non-Hispanic White | | | | | | |
| Non-Hispanic Black | | | -3.17* | 1.26 | -2.01* | 0.88 |
| Hispanic | | | -3.44** | 0.98 | -2.88** | 0.74 |
| Other | | | -0.60 | 1.46 | -1.86 | 1.10 |
| Family poverty | | | | | | |
| <50% | | | -0.01 | 1.29 | 0.34 | 0.58 |
| 51%-100% | | | -0.08 | 1.02 | -0.16 | 0.81 |

| | | | | |
|---|---|---|---|---|
| 101%-200% | | | | |
| >200% | -2.50* | 0.99 | -0.41 | 1.0 |
| Total behavior (Baseline) | | | 0.66** | 0.03 |

*p<.05, **p<.01

*Note*. Model 1 regresses outcome on intelligence at baseline, Model 2 adds covariates, and and Model 3 regresses outcome at Time 2 onto outcome at Time 1, IQ indice, and covariates. + Models with the Passage Comprehension subtest have a sample size of n = 1566 because the subtest was not administered to participants age 12+.
